# Molecular basis for the interactions of eIF2β with eIF5, eIF2B, and 5MP1 and their regulation by CK2

**DOI:** 10.1101/2024.04.25.591181

**Authors:** Paul A. Wagner, Meimei Song, Ralf Ficner, Bernhard Kuhle, Assen Marintchev

## Abstract

The heterotrimeric GTPase eukaryotic translation initiation factor 2 (eIF2) delivers the initiator Met-tRNA_i_ to the ribosomal translation preinitiation complex (PIC). eIF2β has three lysine-rich repeats (K-boxes), important for binding to the GTPase-activating protein eIF5, the guanine nucleotide exchange factor eIF2B, and the regulator eIF5-mimic protein (5MP). Here, we combine X-ray crystallography with NMR to understand the molecular basis and dynamics of these interactions. The crystal structure of yeast eIF5-CTD in complex with eIF2β K-box 3 reveals an extended binding site on eIF2β, far beyond the K-box. We show that eIF2β contains three distinct binding sites, centered on each of the K-boxes, and human eIF5, eIF2Bε, and 5MP1 can bind to all three sites, while reducing each other’s affinities. Our results reveal how eIF2B speeds up the dissociation of eIF5 from eIF2-GDP to promote nucleotide exchange; and how 5MP1 can destabilize eIF5 binding to eIF2 and the PIC, to promote stringent start codon selection. All these affinities are increased by CK2 phosphomimetic mutations, highlighting the role of CK2 in both remodeling and stabilizing the translation apparatus.

## Introduction

Translation initiation in eukaryotes relies on a complex network of interactions that are continuously reorganized throughout the process. As more information becomes available about the structure of the ribosomal preinitiation complex (PIC) at various points in translation initiation, new questions arise about which interactions occur when, their roles, and regulation. The eukaryotic translation initiation factor 2 (eIF2) is a heterotrimeric GTPase playing a central role in the regulation of translation initiation and the integrated stress response (ISR). In its active GTP-bound form, eIF2 binds the initiator Met-tRNA_i_ to form the eIF2-GTP•Met-tRNA_i_ ternary complex (TC). The TC binds to the PIC, either directly or as part of the multifactor complex (MFC), which also includes eIFs 1, 3, and 5. eIF5 is the GTPase-activating protein (GAP) for eIF2. eIF1 serves as a gate-keeper promoting stringency of start codon selection. eIF3 is a large protein complex with multiple roles in translation initiation. Mammalian eIF3 has 13 subunits, while *Saccharomyces cerevisiae (S. cerevisiae)* eIF3 has six (five core subunits and the sub-stoichiometric HCR1/eIF3j). Most of eIF3 resides on the back, solvent-exposed surface of the 40S ribosomal subunit, while some of its subunits reach the interface with the 60S ribosomal subunit. One of them, the N-terminus of eIF3c wraps around the 40S and interacts with eIFs 1, 2, and 5. The interactions among the MFC components are not fully understood but it has become clear that they are remodeled upon binding to the PIC, where the MFC components also interact with the 40S ribosomal subunit, the mRNA, and eIF1A. The complex is further continually remodeled at every step of translation initiation, including displacement of eIF1 from the ribosome by the eIF5 N-terminal domain (NTD) upon start codon selection to form the so-called “closed” PIC, arrested at the start codon (Brito Querido et al., 2024, Dever et al., 2023, Jackson et al., 2010, Marintchev & Wagner, 2004, Petrychenko et al., 2024, Weisser & Ban, 2019).

Following GTP hydrolysis and the release of eIF2-GDP from the PIC, its reactivation to eIF2-GTP and regeneration of the TC require the guanine nucleotide exchange factor (GEF) eIF2B (a decamer with two copies of five different subunits), which promotes nucleotide exchange and subsequent Met-tRNA_i_ binding to eIF2-GTP. At least in *S. cerevisiae*, eIF2-GDP is released from the PIC in a stable complex with eIF5, which serves as GDP dissociation inhibitor (GDI). eIF2B has to displace eIF5 before it can promote GDP dissociation from eIF2, thus acting as GDI dissociating factor (GDF) (Jennings & Pavitt, 2010, Jennings et al., 2013, Singh et al., 2006). As a result, eIF2 is channeled from the PIC, through eIF5, to eIF2B (Jennings & Pavitt, 2010, Jennings et al., 2013, Singh et al., 2006), and could also be directly transferred from eIF2B back to eIF5 and the MFC at the end of the cycle (Bogorad et al., 2018).

eIF2B is one of the main targets in the regulation of protein synthesis in the cell (reviewed in (Bogorad et al., 2018, Brito Querido et al., 2024, Hinnebusch, 2014, Marintchev & Ito, 2020). The activity of eIF2B is regulated by phosphorylation of its substrate eIF2, by binding of nucleotides and cofactors to eIF2B, and by phosphorylation of eIF2B itself by at least two kinases: casein kinase 2 (CK2) stimulates eIF2B activity, while GSK-3β inhibits it. In human, several kinases phosphorylate eIF2α at Serine 51 (S51) in response to various types of stress, including viral infection (PKR), unfolded proteins in the ER (PERK), amino acid starvation (GCN2), and heme deficiency (HRI), in what is collectively known as the Integrated Stress Response (ISR). Phosphorylated eIF2-GDP (eIF2(α-P)-GDP) is a competitive inhibitor of eIF2B. Inhibition of eIF2B activity causes downregulation of global protein synthesis and triggers the ISR by inducing the production of a set of transcription factors and other proteins. The result is the activation of both pro-apoptotic and pro-survival pathways aimed at restoring homeostasis. Since the stressors themselves can cause cell death, either an insufficient or overly aggressive stress response can lead to apoptosis, the latter being the desired outcome in the case of viral infection. Dysregulated ISR is a causative factor in the pathology of a number of neurodegenerative disorders, including Alzheimer’s disease. Mutations in all five subunits of eIF2B cause the genetic disease leukoencephalopathy with vanishing white matter (VWM) (reviewed in (Bogorad et al., 2018, Dever et al., 2007, Hinnebusch, 2005, Hinnebusch et al., 2007, Marintchev & Ito, 2020, Marintchev & Wagner, 2004, Ron & Harding, 2007, Wek et al., 2006).

The eIF5 mimic protein (5MP), also known as BZW, is a regulator that competes for binding to eIF2 with eIF5, leading to increased stringency of start codon selection, and with eIF2B, leading to modulation of nucleotide exchange on eIF2 (Hiraishi et al., 2014, Kozel et al., 2016, Singh et al., 2021, Tang et al., 2017). 5MP also binds to at least one of the other eIF5 partners, eIF3c. While there are two 5MP isoforms in human, 5MP1/BZW2 and 5MP2/BZW1, most of the studies have been performed on 5MP1 (Hiraishi et al., 2014, Kozel et al., 2016, Singh et al., 2021, Tang et al., 2017).

The C-terminal domains (CTD) of eIF5, eIF2Bε, and 5MP are homologous to each other and mediate binding to the intrinsically disordered N-terminal tail of the β subunit of eIF2 (eIF2β- NTT). The interaction is mediated by two acidic/aromatic (AA) boxes in the CTD of eIF5, eIF2Bε, and 5MP and three lysine-rich segments (K-boxes) in eIF2β-NTT (Asano et al., 1999, Das et al., 1997). The contact interfaces of eIF5-CTD with eIF1, 1A, and possibly also 3c, appear to overlap with the eIF2β-NTT binding surface; therefore, eIF2β-NTT is presumed to be at least partially displaced from eIF5 within the PIC (Luna et al., 2013, Luna et al., 2012, Obayashi et al., 2017). eIF5 and eIF2Bε also interact with eIF2γ, with eIF2Bε having two binding sites, one in the NTD and another in the CTD (Alone & Dever, 2006, Kashiwagi et al., 2019, Kenner et al., 2019). While 5MP likely also has additional binding sites (Singh et al., 2021), interactions between 5MP and eIF2 outside the eIF2β-NTT have not yet been reported.

Mammalian eIF5, eIF2Bε, and both 5MP isoforms all have conserved Casein Kinase 2 (CK2) phosphorylation sites in AA-box 2. CK2 phosphorylation of eIF5 leads to stimulation of protein synthesis and cell proliferation (Homma et al., 2005). Phosphorylation of eIF2Bε stimulates eIF2B activity (Singh et al., 1996). The effect of phosphorylation of 5MP1 or 5MP2 has not been investigated. CK2 has hundreds of substrates, including many eIFs, and regulates protein synthesis in a number of ways (Borgo et al., 2017, Franchin et al., 2018, Gandin et al., 2016, Homma et al., 2005, Llorens et al., 2003, Paytubi et al., 2008, Turowec et al., 2010). CK2, together with mTOR, also phosphorylates eIF2β, which helps recruit the PP1 phosphatase that dephosphorylates eIF2α and suppresses the ISR (Gandin et al., 2016). While CK2 is constitutively active in most cells, it is inactivated during chronic stress, leading to dephosphorylation of its substrates and translational reprogramming of the cell (Gandin et al., 2016, Lamper et al., 2020, Lee et al., 2016, Mukhopadhyay et al., 2023).

Despite their importance for translation initiation, the molecular basis for the interactions of eIF2β-NTT with eIF5, eIF2Bε and 5MP, their dynamics and regulation remain poorly understood. The presence of three K-boxes suggests that there could be three binding sites in eIF2β, but this possibility has received little attention because of the 1:1 stoichiometry of the relevant complexes. The first article describing the role of the K-boxes, from the Maitra lab, claimed based on co-IP that a small segment corresponding to K-box 2 is necessary and sufficient for binding to mammalian eIF5 (Das et al., 1997). Asano and co-authors reported the importance of the K-boxes in *S. cerevisiae* eIF2β for binding to both eIF5 and eIF2Bε. While they found that the effects of the K-boxes are largely additive, they concluded based on co-IP that binding of eIF5 and eIF2B is mutually exclusive (Asano et al., 1999). Later, the Asano group found that eIF5 and eIF2B can in fact bind to eIF2 simultaneously but maintained that binding to eIF2β is mutually exclusive and the simultaneous binding is due to additional interactions elsewhere on eIF2 (Singh et al., 2006). Subsequent publications from a number of labs, including ours, as well as reports on 5MP from the Asano lab, also maintained the notion of competition for eIF2β binding (see e.g., (Bogorad et al., 2018, Hiraishi et al., 2014, Jennings et al., 2016, Jennings & Pavitt, 2010, Jennings et al., 2013, Kozel et al., 2016, Luna et al., 2013, Luna et al., 2012, Obayashi et al., 2017, Sato et al., 2019, Singh et al., 2011, Tang et al., 2017). We recently reported that human eIF2β-NTT can bind at least two, and most likely three eIF5- CTD molecules simultaneously (Paul et al., 2022), and that phosphomimetic mutations in eIF5- CTD, mimicking CK2 phosphorylation, increase its affinity for eIF2β (Paul et al., 2022) and eIF1A (Gamble et al., 2022).

Here, we report the crystal structure of the complex of *S. cerevisiae* eIF5-CTD with K-box 3 of eIF2β (eIF2β-K3), which shows an extended binding site on eIF2β that involves segments on both sides of the actual K-box. We go on to show that this binding mode is conserved with human eIF5, as well as eIF2Bε and 5MP1, and that all three proteins can bind to each of the three K-boxes independently and with similar affinities. Finally, phosphomimetic mutations in all three proteins differentially increase their affinities for each K-box and the intact eIF2β-NTT by up to an order of magnitude. Taken together, our results provide both a structural snapshot of the eIF5-eIF2β interactions as well as a plausible model for the dynamic reorganization and regulation of eIF2β-NTT interactions during the translation initiation cycle.

## Results

### Crystal structure of eIF2β-NTT bound to eIF5-CTD

To gain insight into the structural basis for eIF2β-eIF5 interactions, we determined the X-ray crystal structure of *S. cerevisiae* eIF5-CTD bound to the eIF2β-NTT. Among different combinations of eIF5 and eIF2β constructs used for initial crystallization trials, only the complex between eIF5_201-405_ and eIF2β_39-106_, which contains K2 and K3 and flanking regions, produced well-diffracting single crystals in space group P3_2_12 that were suitable for structure determination.

The final atomic model of the eIF5-eIF2β complex, refined at a resolution of 2.0 Å, contains two copies of eIF5_201-405_ per asymmetric unit (AU), each including residues 201 to 399 and additional vector-encoded residues at the N-terminus (_-5_PGLGS_-1_) (details of data collection and refinement statistics are summarized in **Table 1**). Residues 241-399 of both molecules adopt a compact fold composed of four antiparallel helical repeats (R_I_-R_IV_) that is very similar to the previously reported yeast eIF5-CTD structure (Wei et al., 2006): r.m.s.d of 0.8 Å over 133 C_α_ atoms (**Fig 1A**). The N-terminal extension (amino acids 201-240), which is part of the linker between the N- and C-terminal domains of full-length eIF5, folds into two additional α-helices (α1 and α2) that protrude away from the globular CTD and form extensive contacts with symmetry related molecules in the crystal packing. Interestingly, the DWEAR motif, a highly conserved and functionally important region of eIF5, forms the loop between helices α1 and α2 and is inserted into a hydrophobic cleft between α-helices α3, α5, and α8 of another eIF5-CTD molecule in the crystal packing (**Fig EV1**). Notably, although this interaction occurs *in trans*, likely as a crystallographic artifact (**Fig EV1**), a similar interaction could be formed *in cis*, which would allow the DWEAR-motif to associate with the CTD. Such intramolecular contacts between the DWEAR motif and the CTD are supported by NMR chemical shift perturbation (CSP) and deletion analysis (Paul et al., 2022).

**Figure 1.**
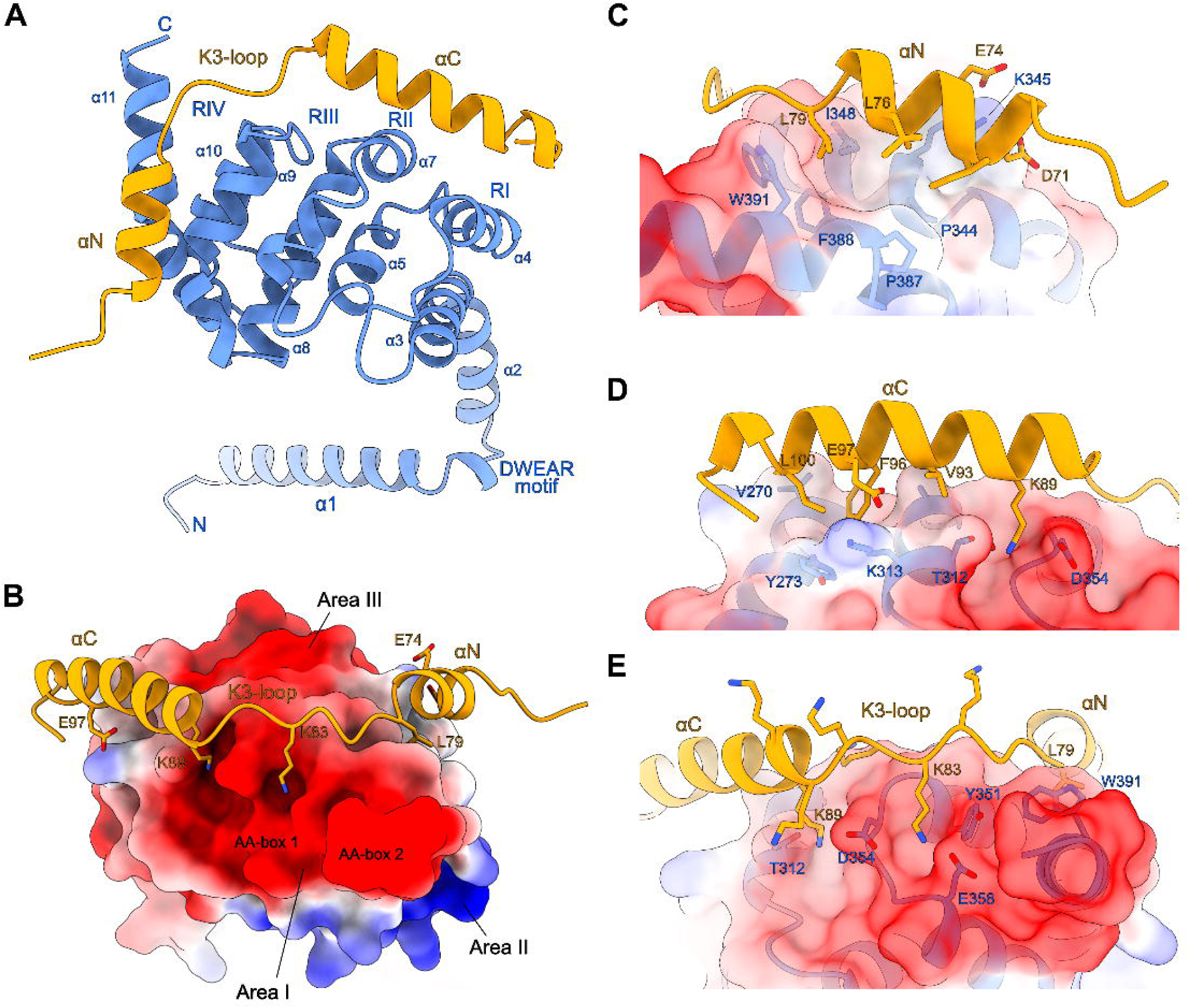
Crystal Structure of the S. cerevisiae eIF5-CTD•eIF2β-K23 complex. (A) Crystal structure of the complex between eIF5-CTD (blue) and eIF2β-K23 (yellow). Residues 39-65 of eIF2β, including K2, are not resolved in the electron density, and are therefore not included in the model. The second eIF5-CTD molecule in the asymmetric unit is not shown. (B) View of the eIF5-CTD•eIF2β-K23 complex centered on the loop containing K3. The eIF5-CTD is shown in surface representation with electrostatic surface charge distribution. (C) Details of the interaction interface between the αN helix of eIF2β-K3 and the surface of eIF5-CTD. (D) Details of the interactions of the αC helix of eIF2β-K3 with eIF5-CTD. (E) Details of the interactions of the K3-loop of eIF2β with eIF5-CTD.

**Table 1.** Data collection and refinement statistics for the structure of the *S. cerevisiae* eIF5_201-405_•eIF2β_39-106_ complex.

| Crystallization |  |
| --- | --- |
| Condition | 0.8 M (NH<sub>4</sub>)SO<sub>4</sub>, 0.3 M LiSO<sub>4</sub> |
| Temperature (°C) | 10 |
| Data Collection |  |
| Space Group | P3<sub>2</sub>12 |
| Unit Cell | a = 74.3 Å<br>b = 74.3 Å<br>c = 124.8 Å |
|  | α = 90°<br>β = 90°<br>γ = 120° |
| Resolution (Å) | 2.0 (2.2-2.0) |
| Observed reflections | 359597 (87743) |
| Unique reflections | 48223 (11831) |
| Redundancy | 7.45 |
| Completeness (%) | 99.9 (99.8) |
| /σ | 22.3 (3.9) |
| R<sub>merge</sub> (%) | 8.9 (42.8) |
| Refinement |  |
| R<sub>work</sub>/R<sub>free</sub> (%) | 20.7/22.8 |
| Wilson B-factor (Å<sup>2</sup>) | 30.4 |
| Total number of atoms | 3824 |
| Average B-factor, all atoms (Å<sup>2</sup>) | 42.0 |
| Rmsd from Strd. Stereochemistry |  |
| Bond length (Å) | 0.015 |
| Bond angles (°) | 1.65 |
| Ramachandran Plot Statistics |  |
| Most favored (%) | 98.9 |
| Allowed regions (%) | 1.1 |
| Disallowed regions (%) | 0 |
| Values in parentheses refer to the highest resolution shell.<br>R<sub>work</sub> and R<sub>free</sub> factors are calculated using the formula $R = \frac{\sum_{hkl} \vert F(\text{obs})_{hkl}\vert - \vert F(\text{calc})_{hkl}\vert }{\sum_{hkl} \vert F(\text{obs})_{hkl}\vert }$, where F(obs)<sub>hkl</sub> and F(calc)<sub>hkl</sub> are observed and measured structure factors, respectively. R<sub>work</sub> and R<sub>free</sub> differ in the set of reflections they are calculated from: R<sub>free</sub> is calculated for the test set, whereas R<sub>work</sub> is calculated for the working set. |  |

A single eIF2β molecule is bound by only one of the two eIF5-CTDs in the AU. This eIF2β fragment includes residues 66-105 and forms two amphipathic α-helices (αN and αC) flanking an unstructured loop containing K-box 3 (**Fig 1A,B**). Residues 39-65 of eIF2β, including K2, are not resolved and thus appear to be flexible. Helices αN and αC are both well-defined in the electron density and form extensive hydrophobic as well as polar interactions with eIF5.

Helix αN (residues 69-79) lies parallel to eIF5 helices α9 and α11 with which it forms a mostly hydrophobic interface (**Fig 1C**). Helix αC (residues 89-103) binds on top of and orthogonal to eIF5 helices α4 and α7, with Phe96 and Leu100 inserted into a hydrophobic pocket on eIF5 and eIF2β-Glu97 forming a salt bridge with eIF5-Lys313 (**Fig 1D**). Between helices αN and αC lies the K3-loop, which is positioned in the immediate vicinity to the negatively charged AA-boxes of eIF5 (area I). Notably, only two of the seven lysine residues, Lys83 and Lys89, form direct interactions with AA-box 1 (**Fig 1E**). The remaining lysine residues of K-box 3 are poorly resolved in the electron density, indicating that they do not participate in stable interactions with eIF5 under the given conditions.

Taken together, the eIF5-bound eIF2β-NTT adopts an elongated conformation, containing secondary, but no tertiary structural features. This allows eIF2β to form an extended interaction interface with eIF5, centered on the K-box and wrapping half around the eIF5-CTD (**Fig 1**). In the following text, ‘K-box’ will refer to this extended binding site, including the adjoining segments αN and αC. eIF2β thereby contacts only the periphery of the negatively charged area I of eIF5, with αN and αC lying between the negatively charged areas I and III and the positively charged area II (**Fig 1B**). Most of area I remains solvent-exposed in the complex, including the highly negatively charged C-terminus (residues 393-405), Glu358, and Glu359, which are critical for high affinity binding of eIF2β-NTT (Das & Maitra, 2000, Yamamoto et al., 2005).

### Human eIF5-CTD, eIF2B**ε**-CTD, and 5MP1-CTD form similar complexes with each of the three eIF2**β**-NTT K-boxes

Multiple-sequence alignment of eIF2β-NTT sequences shows that the segments surrounding the K-boxes are conserved both among species and among the three K-boxes, enriched in acidic and hydrophobic amino acids (**Fig 2A**). If the binding sites centered around K1 and K2 are similar in size to that observed with K3 in the crystal structure (**Fig 1**), then in *S. cerevisiae* eIF2β-NTT, the three binding sites are immediately adjacent to each other, with helix αN of one binding site starting where helix αC of the previous binding site ends (**Fig 2A**). In contrast, in human eIF2β-NTT, there are about 30 amino acids between the predicted K1 and K2 binding site, and about 10 amino acids between K2 and K3 (**Fig 2A**). The spacer between the end of helix αC after K3 and the eIF2γ-binding helix appears to be about 20 amino acids in both human and *S. cerevisiae* eIF2β-NTT.

**Figure 2.**
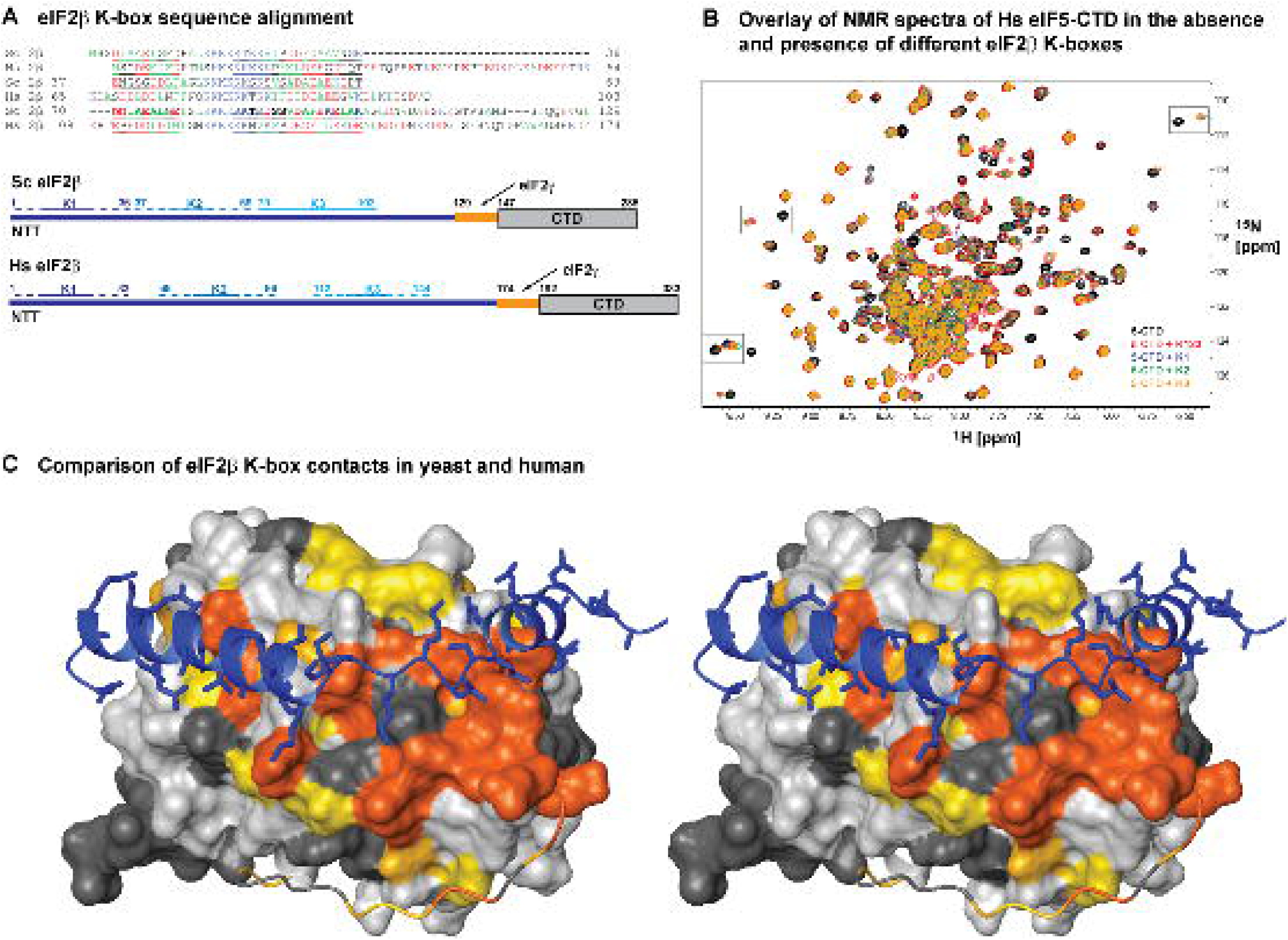
**eIF5-CTD forms similar complexes with each of the three K-boxes of eIF2**β. (A) Sequence alignment of the K-boxes and surrounding sequences in *S. cerevisiae* and human eIF2β- NTT, (labeled “Sc 2β” and “Hs 2β”, respectively). The entire 2β-NTT sequences are shown, up to the start of the eIF2γ-binding helix. Residue numbers are shown at the beginning and end of each line. Positively charged amino acids are colored in blue; negatively charged amino acids are colored in red, and hydrophobic amino acids are colored in green. Helices αN and αC from the *S. cerevisiae* eIF5- CTD•eIF2β-K3 complex (Fig 1) are bold and underlined. The corresponding sequence segments in the other K-boxes in *S. cerevisiae* and human eIF2β are underlined. Note that if the K1 and K2 in *S. cerevisiae* form helices of the same length as those observed for K3 (Fig 1), then there are no linker amino acids between K1 and K2, and between K2 and K3 in *S. cerevisiae* eIF2β. Diagrams of *S. cerevisiae* and human eIF2β are shown below the alignment. Folded domains are shown as boxes. Intrinsically disordered segments are shown as lines. K-boxes K1, K2, and K3 are marked and colored dark blue, blue, and light blue, respectively. The K3 segment of *S. cerevisiae* eIF2β visible in the crystal structure, including the αN and αC helices (Fig 1) is marked with a solid line. The corresponding segments surrounding the other K-boxes, corresponding to the αN and αC helices are marked with dashed lines. (B) NMR spectra overlay of ^15^N/^2^H-labeled human eIF5-CTD, free (black) and in the presence of K123 (red), K1 (blue), K2 (green), and K3 (orange). Examples of peaks that are in similar positions in all complexes (top two boxes), or different positions in the different complexes (bottom left) are boxed. (C) Comparison of eIF2β K-box contacts in yeast and human, in cross-eyed stereo. The eIF2β-K3 segment from the crystal structure of the *S. cerevisiae* eIF5-CTD•eIF2β-K3 complex (Fig 1) is overlayed on the structure of human eIF5-CTD (2iu1.pdb(Bieniossek et al., 2006)). The overlay was obtained by aligning the structures of human and yeast eIF5-CTD. eIF2β is shown as a blue ribbon, with sidechains also shown. Human eIF5-CTD is shown in surface representation colored by CSP effects for K123 binding: from dark orange (large effects) to yellow (small effects). No significant changes are light grey; amino-acids that could not be analyzed are in dark grey. eIF5-CTT is disordered in solution(Paul et al., 2022) and shown as a ribbon.

We used NMR Chemical Shift Perturbation (CSP) assay to compare the contact surfaces of human eIF5-CTD for each of the three K-boxes and the complete eIF2β-NTT. All further experiments described below use human proteins (constructs used in this work are shown in **Fig EV2A**). Comparison of the ^15^N-transverse relaxation optimized spectroscopy – heteronuclear single quantum coherence (TROSY-HSQC) NMR spectra of free ^15^N/^2^H-labeled eIF5-CTD and of eIF5-CTD in complex with unlabeled individual K-boxes and eIF2β-NTT shows that the vast majority of the peaks affected by binding are the same in all four complexes, with even the magnitude of the peak movement being similar as well (**Fig 2B and EV2B**). Mapping the affected amino acids on the surface of eIF5-CTD (**Fig 2C**) shows that the contact interface is similar to that observed in the crystal structure of the yeast eIF5-CTD•eIF2β-K3 complex (**Fig 1**), and also extends into the intrinsically disordered eIF5 C-terminal tail (**Fig 2C**). Note that in **Fig 2C**, the position of eIF5-CTT is from the crystal structure of human eIF5-CTD (2iu1.pdb) (Bieniossek et al., 2006). However, we have previously shown that the CTT is intrinsically disordered and remains disordered while contacting eIF2β (Paul et al., 2022). Accordingly, the corresponding region of *S. cerevisiae* eIF5 (residues 400-405), which is not resolved in the crystal structure (**Fig 1**), could also be involved in dynamic interactions with eIF2β. Consistent with the crystal structure of the *S. cerevisiae* complex, mutating hydrophobic residues N-terminal from K1, corresponding to helix αN, or deleting a segment C-terminal from K3, corresponding to the C-terminal portion of helix αC, weakens binding to eIF5-CTD by at least an order of magnitude, based on the concentration of the labeled eIF5-CTD (50 μM) and the unlabeled mutant K-boxes (25 μM) (**Fig EV3A** and **B**, respectively). Therefore, human eIF5- CTD forms very similar complexes with all K-boxes, each likely similar to the complex of *S. cerevisiae* eIF5-CTD with eIF2β-K3 (**Fig 1**).

NMR backbone resonance assignments are not available for eIF2Bε-CTD and 5MP1- CTD, meaning that we do not know which peak in the TROSY-HSQC spectra corresponds to which residue in the protein. However, as with eIF5-CTD, most peaks in the eIF2Bε-CTD and 5MP1-CTD spectra, affected by binding of the three individual K-boxes or the entire eIF2β-NTT, are the same in all four complexes, with even the magnitude of the peak movement being similar (**Fig EV4A,B**). Therefore, as for eIF5-CTD, the same surfaces of eIF2Bε-CTD and 5MP1-CTD are affected in all four complexes. Thus, all three proteins form similar complexes with all three K-boxes, both in isolation and when part of the three-K-box eIF2β-NTT (**Fig 2, EV2, and EV4**).

### Human eIF5-CTD, eIF2B**ε**-CTD, and 5MP1-CTD have distinct binding affinities for eIF2**β**- NTT and individual K-boxes

We used NMR CSP assay titrations to obtain the K_D_s of eIF5-CTD, eIF2Bε-CTD, and 5MP1-CTD binding to each of the three K-boxes and the intact eIF2β-NTT (**Fig 3** and **Table 2**). Binding is in fast to intermediate exchange on the NMR time scale, depending on the degree of change in peak position (**Fig EV4C**). Peaks that move by > 100 Hz (0.2 ppm for ^1^H on a 500 MHz instrument) undergo exchange broadening; therefore, under the experimental conditions (150 mM KCl and 25 LJC), k_off_ is in the order of 100 s^-1^ (reviewed in (Marintchev et al., 2007)). The effect of eIF2β-NTT binding is the weighted average of the binding of the three individual K-boxes. The presence of one average peak for the bound state indicates that each molecule of the labeled protein constantly samples all three K-boxes on a millisecond time scale (**Fig 2B** and **EV4**).

**Figure 3.**
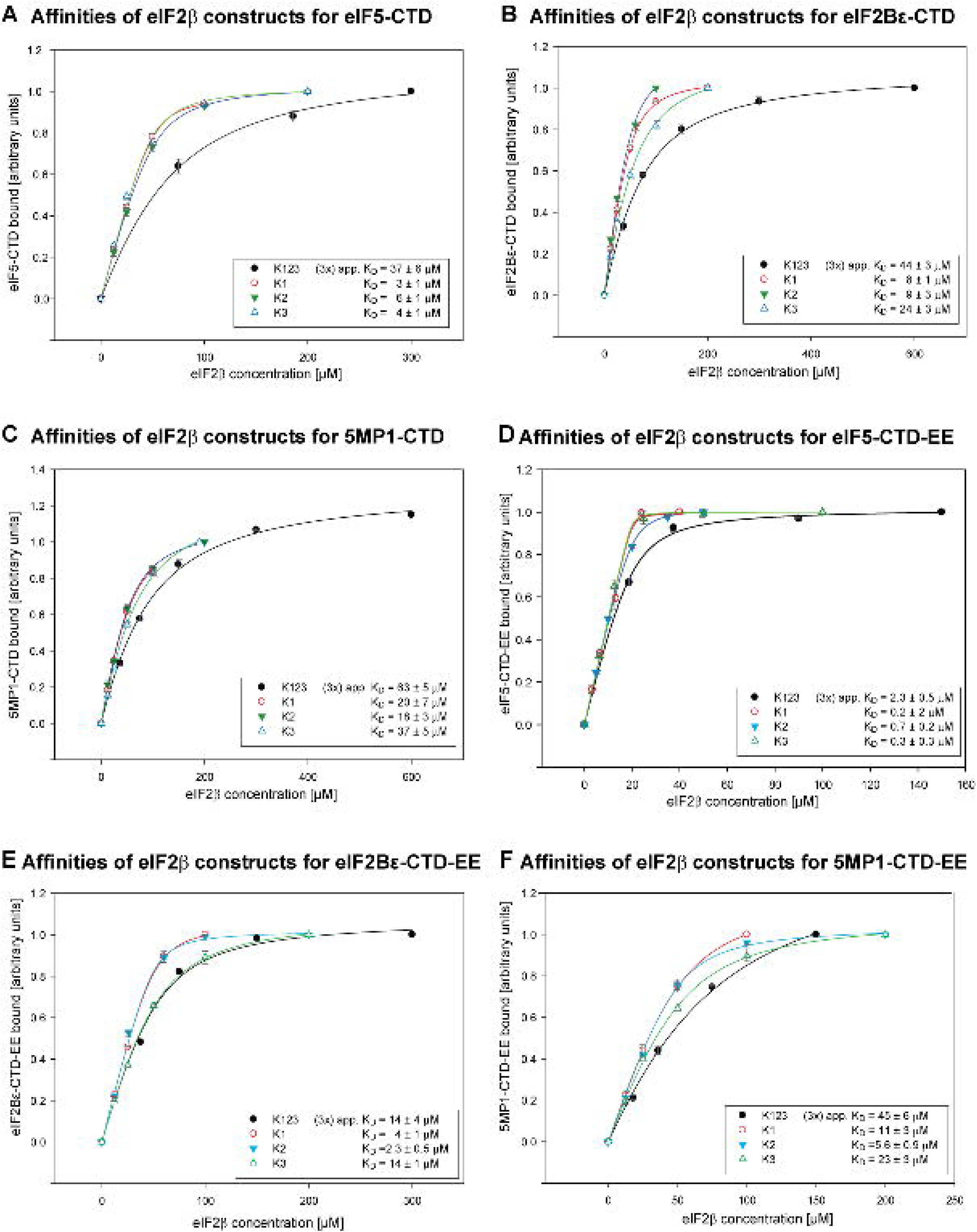
**Binding affinities of WT and phosphomimetic human eIF5-CTD, eIF2B**ε**-CTD, and 5MP1- CTD for eIF2**β**-NTT and its individual K-boxes.** (A) Overlay of titration curves of eIF5-CTD binding to K1, K2, K3, and K123. (B) Overlay of titration curves of eIF2Bε-CTD binding to K1, K2, K3, and K123. (C) Overlay of titration curves of 5MP1-CTD binding to K1, K2, K3, and K123. (D) Overlay of titration curves of eIF5-CTD-EE binding to K1, K2, K3, and K123. (E) Overlay of titration curves of eIF2Bε-CTD-EE binding to K1, K2, K3, and K123. (F) Overlay of titration curves of 5MP1-CTD-EE binding to K1, K2, K3, and K123. Note that the K_D_s for K123 are averages, assuming three binding sites with identical affinities. Standard deviations (SD) are shown as grey bars. The chemical shift change values were normalized to the point with the greatest chemical shift change (assigned a value of 1); therefore, the last point has no SD values. The calculated K_D_s and standard errors (SE) are shown in the insets.

**Table 2.** Binding affinities of WT and phosphomimetic eIF5, eIF2B*ε*, and 5MP1 for the eIF2*β* K-boxes.

| | $K_D$ [μM] | | | |
| --- | --- | --- | --- | --- |
|  | K1 | K2 | K3 | K123 |
| eIF5-CTD | $3 \pm 1$ | $6 \pm 1$ | $4 \pm 1$ | (3x) $37 \pm 6^1$ |
| eIF2Bε-CTD | $8 \pm 1$ | $9 \pm 3$ | $24 \pm 3$ | (3x) $44 \pm 3$ |
| 5MP1-CTD | $20 \pm 7$ | $18 \pm 3$ | $37 \pm 5$ | (3x) $63 \pm 5$ |
| eIF5-CTD_EE | $0.2 \pm 0.2^2$ | $0.7 \pm 0.2^2$ | $0.3 \pm 0.3^2$ | (3x) $2.3 \pm 0.5$ |
| eIF2Bε-CTD_EE | $4 \pm 1$ | $2.3 \pm 0.5$ | $14 \pm 1$ | (3x) $14 \pm 4$ |
| 5MP1-CTD_EE | $11 \pm 3$ | $5.6 \pm 0.9$ | $23 \pm 3$ | (3x) $45 \pm 6$ |

<sup>1</sup> The $K_D$ s for K123 are averages, assuming three binding sites with identical affinities.
<sup>2</sup> $K_D$ values below 1 μM are only approximate because the concentration of the labeled protein (20-50 μM) becomes much larger than the $K_D$ , making accurate fitting difficult.

Because eIF2β-NTT has three non-equivalent binding sites, obtaining all three K_D_s reliably is only possible if they differ by at least 5-fold (and the three binding sites are occupied mostly consecutively). However, at least the average K_D_ (assuming three binding sites with identical affinities) can be obtained. We found that eIF5-CTD binds to all three eIF2β-NTT K-boxes with similar affinities (**Fig 3A**). In contrast, eIF2Bε-CTD has greater affinity for K2 and K1 than for K3 (**Fig 3B**). As a result, eIF5-CTD has a much higher affinity for K3 than eIF2Bε-CTD, whereas their affinities for K1 and K2 are comparable (compare **Fig 3A and B**). 5MP1-CTD binds with similar affinities to all three eIF2β-NTT K-boxes (**Fig 3C**), a few-fold weaker than eIF5-CTD and eIF2Bε-CTD (**Fig 3A,B,C**, **Table 2**). The relative affinities for a given K-box ranged from similar (eIF5-CTD and eIF2Bε-CTD for K2) to different by an order of magnitude (eIF5-CTD and 5MP1-CTD for K3). Overall, K3 shows clear preference for eIF5-CTD, whereas the K_D_s of the proteins for K1 and 2 are much closer (**Fig 3A,B,C**, **Table 2**).

Surprisingly, the affinities for eIF2β-NTT (containing K-boxes 1, 2, and 3) are lower than expected from the K_D_s for the individual K-boxes. For eIF2Bε-CTD and 5MP1, the average K_D_, calculated assuming three binding sites with equal affinities, is not only higher than the average of the K_D_s for the three individual K-boxes, but 2-fold higher than the highest K_D_. For eIF5-CTD, this difference is even greater: the average K_D_ for eIF2β-NTT is 6-fold higher than the highest individual K_D_, that for K2 (**Fig 3A,B,C**, **Table 2**). As a result, the overall affinities of all three proteins for eIF2β-NTT end up being similar to each other (**Table 2**). As described above, the peak positions in the presence of eIF2β-NTT are the weighted average of their positions in the presence of the three individual K-boxes. This weighted average appears similar to the relative affinities for the three K-boxes measured in isolation, both when eIF2β-NTT is in excess and there is on average only one occupied K-box (**Fig 2B** and **EV4A,B**), and when its binding partner is in excess, and all three sites are partially occupied (**Fig EV5A**). Therefore, it appears that simultaneous binding to all three K-boxes is weaker when part of the eIF2β-NTT.

Similarly, NMR spectra of eIF2β-NTT in the presence of equimolar amounts of eIF5- CTD, 5MP1-CTD, and eIF2Bε-CTD showed that the positions of the peaks affected by binding were the weighted average of their positions in the presence of each of the three individual proteins (**Fig EV5A**). Therefore, each K-box binds to each of the proteins on a millisecond time scale. We used size exclusion chromatography (SEC) to visualize directly the formation of a complex of eIF2β-K12 (containing K-boxes 1 and 2) with eIF5-CTD and eIF2Bε-CTD. As expected, when mixed at equimolar concentrations, eIF5-CTD and eIF2Bε-CTD comigrate with eIF2β-K12, which demonstrates that eIF5-CTD and eIF2Bε-CTD can bind simultaneously to different K-boxes on the same eIF2β molecule (**Fig EV5B**).

### Phosphomimetic mutations in human eIF5-CTD, eIF2B**ε**-CTD, and 5MP1-CTD differentially enhance binding to eIF2**β**-NTT and individual K-boxes

Mammalian eIF5-CTD, eIF2Bε-CTD, and 5MP1-CTD all have conserved CK2 phosphorylation sites in their AA-box 2, which is important for eIF2β binding. While phosphorylation of eIF5-CTD (Homma et al., 2005) and eIF2Bε-CTD (Singh et al., 1996) by CK2, and its biological effects have been studied *in vitro*, phosphorylation of the 5MPs has only been observed in high-throughput studies (see e.g. (Mertins et al., 2016, Yi et al., 2014)).

Therefore, we first confirmed that 5MP1-CTD is readily phosphorylated by CK2 *in vitro*. Phosphomimetic mutations mimicking CK2 phosphorylation in all three proteins increased their affinities for eIF2β-NTT and the individual K-boxes (**Fig 3D,E,F**, **Table 2**). However, the degree of increase varied significantly, from 50% (5MP1-CTD-EE with K3) to an order of magnitude (eIF5-CTD-EE with K2). The increase in average affinity for eIF2β-NTT also ranged from 50% (5MP1-CTD-EE) to an order of magnitude (eIF5-CTD-EE) (**Table 2**). The overall pattern observed was that the phosphomimetic mutations in eIF5-CTD increased its affinity to a greater extent than those in eIF2Bε-CTD or 5MP1-CTD. Among the different eIF2β constructs tested, the phosphomimetic mutations increased the affinities for K2 the most (at least three-fold or greater change in K_D_). Notably, the affinity of eIF2Bε-CTD-EE for K123 was equal to its affinity for K3, while still being much weaker than the average of its affinities for the three individual K-boxes. The phosphomimetic mutations changed the relative affinities among the three proteins for a given K-box by up to three-fold, in favor of eIF5-CTD (**Table 2**). Therefore, phosphorylation by CK2 modulates the relative affinities of the individual proteins for eIF2β-NTT, in addition to increasing all their affinities.

## Discussion

In the crystal structure of the *S. cerevisiae* eIF2β-K3•eIF5-CTD complex (**Fig 1**), eIF2β- K3 and the adjoining sequences wrap half-around eIF5-CTD and fold upon binding, forming two helices, αN and αC. The K-box itself forms the loop between the two helices and the N-terminal portion of αC (**Fig 1**). Notably, helix αC ends just above the first of the four antiparallel helical repeats (R_I_) of eIF5-CTD (**Fig 1**), thus supporting the role of eIF2β-NTT in recruiting and positioning the eIF5-CTD and the preceding DWEAR motif relative to the eIF2αβγ complex. The interaction of eIF5-CTD with a portion of the eIF2β-NTT was recently modeled in the context of the 43S PIC from *Trypanosoma cruzi (T. cruzi*). The poor map quality in the corresponding region and the absence of side chain information allowed only a tentative placement of two α- helices, which were interpreted as belonging to K-box 3 of eIF2β (chain **n** in 7ASE.pdb, labeled as “Translation initiation factor, putative”) (Bochler et al., 2020). Notwithstanding the lack of molecular details, this interpretation is confirmed by our crystal structure of the yeast eIF5-CTD - eIF2β complex, which would suggest that the interactions between eIF5-CTD and the eIF2β K-boxes are conserved throughout eukaryotes. Interestingly, K3 is the least conserved K-box in kinetoplastids, with only 2-3 positive charges in *Trypanosoma* eIF2β and no positive charges in *Leishmania* eIF2β (see Fig. S2B in ref. (Bochler et al., 2020)). Thus, should *T. cruzi* eIF5-CTD indeed bind to K3 as proposed in (Bochler et al., 2020), this would further highlight the importance of the regions flanking the K-box itself. Our NMR data (**Fig 2** and **EV4**) show that each of the three K-boxes in human eIF2β forms similar complexes with eIF5-CTD, eIF2Bε- CTD, and 5MP1-CTD, suggesting a common interface similar to that observed in the *S. cerevisiae* eIF2β-K3•eIF5-CTD complex (**Fig 1**).

The affinities of individual proteins to different K-boxes vary by up to an order of magnitude, indicating defined preferences of proteins for one K-box over another, and preferences of K-boxes for one protein over another, respectively. At the same time, most of the individual affinities are high enough and similar enough to be physiologically relevant (**Table 2**). In *S. cerevisiae*, GCN2 phosphorylates eIF2β on Ser80 in K-box 3, which increases its affinity for eIF5, but has no effect on its affinity for eIF2Bε (Dokladal et al., 2021). This finding provides indirect evidence that in *S. cerevisiae*, eIF5 may prefer K3, while eIF2Bε could bind preferentially to K1 or 2, which is consistent with the binding affinities we measured for the human proteins (**Table 2**).

The binding affinities for the intact eIF2β-NTT and individual K-boxes (**Table 2**) indicate that the three K-boxes interfere with each other’s binding, especially to eIF5. While in *S. cerevisiae*, the three binding sites are adjacent to each other, in human eIF2β-NTT, the K-boxes are farther apart (**Fig 2A**); thus, steric interference was not expected *a priori*. It is not clear whether this difference between human and yeast eIF2β has any functional significance.

However, it should be pointed out that in humans, eIF2β-NTT has additional regulatory roles: it has a binding site for raptor (and thus MTORC1) and is phosphorylated by both MTORC1 and CK2, which promotes binding of NCK1, an adaptor of the PP1 phosphatase, leading to eIF2α dephosphorylation and blunting of the ISR (Gandin et al., 2016). It is possible that a protein bound to one K-box and the surrounding segments could transiently contact another K-box.

Such contacts could interfere with a second protein binding to the second K-box. Consistent with this idea, there are negatively charged surfaces in eIF5-CTD that remain unoccupied in the crystal structure of the complex with K3 (**Fig 1**) but display NMR CSP effects (**Fig 2**), indicating that they contact the K-box at least dynamically. It is conceivable that a second K-box could compete for these transient interactions with the “main” K-box. In view of the greater proximity of the individual K-boxes in *S. cerevisiae* (**Fig 2A**), it is possible that the degree of interference between proteins bound simultaneously to adjacent K-boxes could be even greater than observed here for the human proteins. Unfortunately, we were unable to obtain evidence for or against a second K-box simultaneously contacting this surface in eIF5-CTD.

What is the stoichiometry *in vivo*? Extensive proteomics data in paxdb.org (Huang et al., 2023) shows that in human cells, the combined number of eIF5, eIF2B, 5MP1, and 5MP2 molecules is greater than that of eIF2, typically 1.5-fold, and up to twice as large. The same is true in *S. cerevisiae*, which does not have a 5MP homolog, but the combined concentrations of eIF5 and eIF2B are greater than that of eIF2, again about 1.5-fold. Therefore, *in vivo*, the majority of eIF2β-NTT would typically be bound to one or two of these proteins. As to why, the most likely answer is regulation. Specifically, formation of a transient complex, where two proteins are bound simultaneously to eIF2β-NTT, likely allows fast displacement of one protein bound to the trimeric eIF2 by the other. For example, this could be involved in eIF2B displacing eIF5 from eIF2-GDP in order to initiate nucleotide exchange on eIF2 (Jennings & Pavitt, 2010, Jennings et al., 2013, Singh et al., 2006). Such speeding up of the interchange between eIF2- bound proteins would be even more efficient if simultaneous binding of two proteins to eIF2β- NTT weakens their affinities (**Table 2**). Consistent with this idea, mutating any of the three K-boxes in *S. cerevisiae* eIF2β causes a general control derepressed (Gcd^-^) phenotype, indicating lower activity of eIF2B that in turn leads to lower than normal levels of the eIF2-GTP•Met-tRNA_i_ ternary complex (Asano et al., 1999). At the same time, the mutant strains are viable and grow at WT rates in rich medium; and at least K-boxes 1 and 3 alone are sufficient for viability (Asano et al., 1999). Therefore, the presence of multiple K-boxes serves mostly regulatory roles, at least in *S. cerevisiae*.

How is eIF2 handed off between eIF5 and eIF2B, and how do the 5MPs modulate eIF5 and eIF2B binding to eIF2? Both eIF2B and eIF5 contact eIF2 at two or more separate surfaces, yielding a stable overall interaction; and the same is probably the case also for the 5MPs (Singh et al., 2021). Binding to each individual site is weaker, with faster off-rates, but after transient loss of one interaction, the other contact is sufficient to maintain the integrity of the complex and keep the protein in the vicinity of the other site, increasing the rate of re-binding. Our results show that eIF5, eIF2Bε, and 5MP1 bind each of the three K-boxes multiple times per second (**Fig 2B** and **EV4A,B**), and each K-box binds eIF5, eIF2Bε, and 5MP1 multiple times per second (**Fig EV4C**). Therefore, the proteins modulate each other’s binding to eIF2 both by competing for the same K-boxes and by binding simultaneously to different K-boxes. Simultaneous binding to eIF2β-NTT has two complementary effects (**Fig 4**):

- Destabilization of each other’s binding to eIF2β-NTT (**Table 2**), leading to accelerated dissociation (illustrated with red T-shaped lines in the model in **Fig 4**).
- Increased effective concentration in the vicinity of their other binding sites on eIF2.

**Figure 4.**
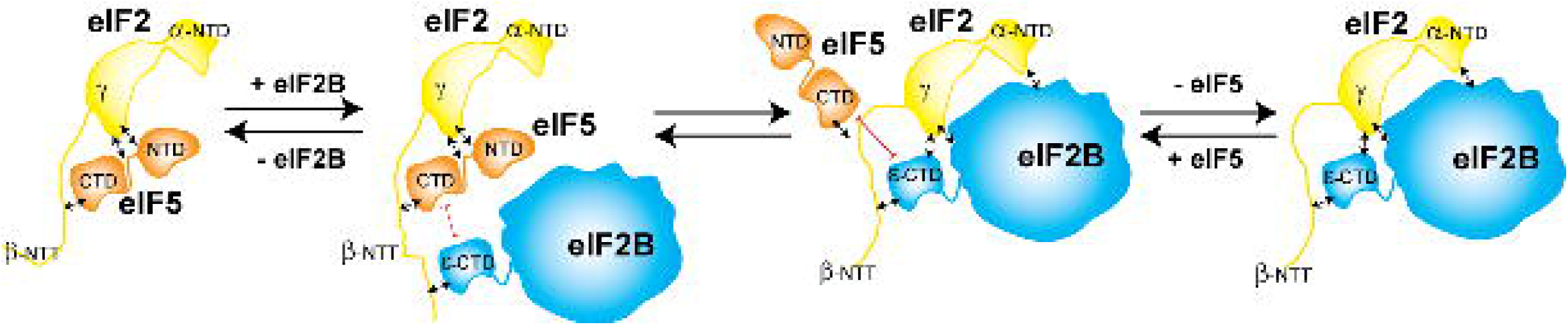
Model for the handoff of eIF2 between eIF5 and eIF2B. Folded domains are shown as shapes. Intrinsically disordered regions are shown as lines. eIF2 is yellow; eIF5 is brown; eIF2B is blue. The shapes of the folded segments of eIF2 and eIF2B, their mutual orientations, and contacts are based on the eIF2B•eIF2 complex structure (6jlw.pdb (Kashiwagi et al., 2019)). For simplicity, only one eIF2 is shown bound to eIF2B. Interactions are shown with double arrows, instead of with physical contacts in the complex, to emphasize their dynamic nature. The mutual destabilization between two proteins simultaneously bound to eIF2β-NTT is illustrated by red T-shaped lines. The same mechanism could work in the opposite direction in the handoff of the TC from eIF2B to eIF5, except there eIF2α-NTD would be interaction with the Met-tRNA_i_, instead of with eIF2B.

For example, eIF2Bε-CTD binding to a different K-box on eIF2β, simultaneously with eIF5-CTD, would increase the probability of eIF2B binding to eIF2γ when eIF5 transiently vacates its binding site there, which in turn would accelerate eIF5 dissociation from eIF2 (**Fig 4**).

All of this applies equally to eIF5 displacing eIF2B, presumably from the TC, if TC dissociation from eIF2B is too slow and/or the equilibrium is shifted too far toward the eIF2B•TC complex (Bogorad et al., 2018). In the eIF2B•TC complex, eIF2α-NTD would likely contact the Met-tRNAi, and not eIF2B, which would presumably weaken the eIF2B – eIF2 interaction, but not the eIF5 – eIF2 interaction. As a result, by simultaneously binding to eIF2, eIF5 and eIF2B weaken each other’s affinities, leading to accelerated dissociation rates and exchange.

Likewise, simultaneous binding would allow the 5MPs to modulate the eIF2 affinity for eIF5 and eIF2B without the need to win a direct competition.

The stringency of start codon selection is controlled by the interplay among eIF1, eIF5, and the 5MPs; and these proteins regulate the translation of their own and each other’s mRNAs through their effects on the stringency (Ivanov et al., 2010, Loughran et al., 2018, Loughran et al., 2012). If the stringency of start codon selection depends on the concentrations of eIF1, eIF5 and the 5MPs, their binding to the PIC must be reversible and in equilibrium with the free proteins (occupancy must be at least slightly below 100%). Our results show how the 5MPs and eIF5 can co-exist on the PIC, weaken each other’s binding affinity for the PIC, and thus lower each other’s occupancy on eIF2 and the PIC. A similar mechanism should be in place for eIF1 and eIF5. Grosely and coauthors recently reported that in an *in vitro* reconstituted system, eIF1 and eIF5 binding to the PIC is indeed dynamic (Grosely et al., 2024). Here, we show that CK2 not only shifts the equilibrium among multiple competing interactions, but also stabilizes all of them (**Table 2**), potentially stabilizing the PIC and making the entire process more efficient. As described above, the stringency of start codon selection is controlled by the interplay among eIF1, eIF5, and the 5MPs (Ivanov et al., 2010, Loughran et al., 2018, Loughran et al., 2012). Therefore, translation initiation at suboptimal start codons will likely also be modulated by eIF5 and 5MP phosphorylation. Dephosphorylation of eIF5 under chronic stress would lower its affinities for eIF1A (Gamble et al., 2022) and eIF2; and the resulting destabilization of the PICs would coincide with a drop in overall protein synthesis, and thus cellular concentrations of ribosomes and eIFs, which are proportional to the rates of protein synthesis (Milo et al., 2010). This raises the possibility for the formation of PICs with alternative properties under these conditions. In view of all these considerations, it would be very intriguing to elucidate how the translation apparatus operates under chronic stress.

The results presented here indicate that the differences in translation under various conditions, e.g., under chronic stress vs. actively dividing cells, may not be as simple as having different numbers of ribosomes and translation factors, but likely include kinetic and possibly even structural differences in the translational apparatus. This reinforces the need to always be mindful of the posttranslational modifications present under the specific conditions studied, especially when performing quantitative analysis or modeling.

## Materials and Methods

### Protein expression and purification

All *S. cerevisiae* constructs were expressed in *E. coli* BL21(DE3) Rosetta II cells (Novagene). Cells transformed with the respective plasmid were grown in 2xYT (containing the appropriate antibiotics) at 37 °C with shaking at 220 rpm until they reached an OD_600_ of ∼0.8 and were subsequently transferred to 16 °C. After allowing the cultures to cool down for 20 min, the expression was induced by the addition of isopropyl-β-D-thiogalactopyranosid (IPTG) to a final concentration of 0.5 mM. The cells were harvested after 16 hours at 16 °C. All human constructs were expressed in *E. coli* BL21(DE3) (Novagene), grown in LB at 37 °C. Expression was induced at 20 °C O/N with 1 mM IPTG. ^15^N/^2^H-labeled proteins were expressed in minimal medium in ^2^H_2_O, with ^15^N-NH_4_Cl as the only nitrogen source, using the same procedure. ^15^N^2^H-labeled eIF2Bε-CTD-EE had poor solubility. Therefore, ^15^N-labeled eIF2Bε-CTD-EE was used, instead, expressed in minimal medium with ^15^N-NH_4_Cl as the only nitrogen source, using the same procedure.

All yeast eIF2β variants were cloned into the pGEX-6P1 vector and expressed as N-terminal GST-fusion proteins. Cells were resuspended in lysis buffer (500 mM NaCl, 20 mM HEPES (pH 7.5), 5% glycerol, 0.1 mM EDTA, 4 mM β-mercaptoethanol (BME)) supplied with a mixture of protease inhibitors including aprotinin, leupeptin, pepstatin (ALP), and PMSF. The cells were lysed in a microfluidizer (Microfluidics, USA) and cell debris was removed by centrifugation for 30 min at 30,000xg. After centrifugation, the supernatant was loaded onto a GSTrap column (GE Healthcare) equilibrated in lysis buffer. The column was washed with 2 column volumes of lysis buffer and bound fusion protein was eluted lysis buffer containing 30 mM of reduced glutathione. Fractions containing target protein were pooled and incubated over night at 4°C with PreScission protease (GE Healthcare) at a ratio of 1:100 (w/w) of protease to fusion protein to remove the GST-tag. After a desalting step in 200 mM NaCl, 20 mM HEPES pH 7.5, 5% glycerol, 4 mM BME, the cleaved GST, PreScission protease, and uncleaved protein were removed by a second GSH-Sepharose step. The flow-through containing the cleaved eIF2β was pooled and applied to a Superdex S200 column (GE Healthcare) equilibrated in a buffer containing 150 mM KCl, 10 mM HEPES (pH 7.5), 5% glycerol, 2 mM DTT.

All yeast eIF5-CTD constructs were cloned into pGEX-6P1 for expression as N-terminal GST-fusions and purified similar to eIF2β with minor modifications. Following lysis and sample application in lysis buffer, the GSTrap column was equilibrated in low salt buffer (100 mM NaCl, 10 mM HEPES pH 8, 5% glycerol, 4 mM BME), followed by elution in low salt buffer containing 30 mM reduced glutathione. After removal of the GST-tag by PreScission protease treatment, the protein was loaded onto a Source 30Q column (GE Healthcare) equilibrated in low salt buffer. Bound eIF5-CTD was eluted with a linear gradient into high salt buffer (1 M NaCl, 10 mM HEPES pH 7.5, 5% glycerol, 4 mM BME). Fractions containing the target protein were pooled and loaded onto a Superdex S75 column (GE Healthcare) equilibrated in buffer containing 150 mM KCl, 10 mM HEPES pH 7.5, 5% glycerol, 2 mM DTT. Fractions containing pure target protein were pooled, concentrated, flash-frozen in liquid nitrogen, and stored at-80 °C.

Expression and purification of recombinant His_6_-tagged human eIF2β-NTT (residues 1- 191) (Luna et al., 2012), eIF5-CTD_-CTT_ (residues 232-431) and its phosphomimetic mutant (Paul et al., 2022) was as described previously. All other human constructs were cloned in pET21a with an N-terminal GB1 tag, a His_6_-tag and a TEV protease cleavage site. Phosphomimetic point mutations were generated by site-directed mutagenesis. The proteins were purified on a TALON Cell-Thru His-tag affinity column (Clontech) in a buffer containing 10 mM Na Phosphate (pH 7.0), 300 mM KCl, 7 mM BME, and 0.1 mM AEBSF, followed by ion exchange chromatography on a Uno Q column, using a 100 mM to 1 M NaCl gradient. Where necessary, the GB1 tag was cleaved using TEV protease. Ion exchange chromatography on a Uno Q column was used to remove the GB1 tag. Proteins were exchanged into buffer containing 10 mM Na Phosphate (pH 7.0), 150 mM KCl, 1 mM EDTA, 0.02% NaN_3_, 1 mM DTT, and 0.1 mM AEBSF.

### Reconstitution, crystallization, and structure determination of the eIF5-eIF2**β** complex

The individual yeast proteins, eIF2β_39-106_ and eIF5_201-405_, were purified as described above. The complex was reconstituted in binding buffer (150 mM KCl, 10 mM HEPES pH 7.5, 5% glycerol, 2 mM DTT) by mixing eIF5_201-405_ with a 1.5-fold molar excess of eIF2β_39-106_. After incubation for 30 min at 20 °C, the complex was separated from excess eIF2β_39-106_ by size exclusion chromatography on a Superdex S75 equilibrated in binding buffer. The purified complex was concentrated to ∼20 mg/ml and directly used for crystallization trials.

Crystallization trials were performed using the sitting drop vapor diffusion method. Initial crystals of the eIF5_201-405_/eIF2β_39-106_complex grew within one day at 20 °C in a condition containing 0.4 M (NH_4_)_2_SO_4_ and 0.05 M Li_2_SO_4_. After optimization, high-quality crystals that were used for structure determination were obtained with 12 mg/ml protein in 0.4 M (NH_4_)_2_SO_4_ and 0.08 M Li_2_SO_4_ at 10 °C.

X-ray diffraction data were collected at BL 14.1 (HZB, BESSY, Berlin) (Mueller et al., 2012). The obtained data were processed in space group P3_2_12 using XDS (Kabsch, 2010) and scaled to a final resolution of 2.0 Å. The phase problem was solved by molecular replacement using the program PHASER (McCoy et al., 2007) with the atomic coordinates of yeast eIF5_241-396_ (PDB 2FUL) as search model. The initial structural model comprised two eIF5_241-396_ molecules per asymmetric unit. Missing regions of the eIF5 construct (residues 201-240) and eIF2β were built manually in Coot (Emsley et al., 2010). The final model was obtained gradually by iterative rounds of refinement in PHENIX (Adams et al., 2010), followed by manual model building.

### Nuclear Magnetic Resonance (NMR)

NMR experiments were performed in buffer containing 10 mM Na Phosphate, pH 7.0, 150 mM KCl, 1 mM EDTA, 0.02% NaN_3_, 1 mM DTT and 0.1 mM AEBSF, with 5% ^2^H_2_O. NMR data were collected on a 500 MHz Bruker spectrometer (Boston University School of Medicine) equipped with a cryoprobe. NMR resonance assignments for eIF5-CTD were available (Paul et al., 2022).

### NMR Chemical Shift Perturbation (CSP) assay

The CSP assay was performed using Heteronuclear single-quantum coherence (HSQC) experiments on ^15^N-labeled proteins. ^15^N Transverse relaxation optimized spectroscopy (TROSY) HSQC experiments were run on ^15^N/^2^H-labeled proteins. Chemical shift changes were calculated according to the formula δ = ((δ_H_)^2^ + (δ_N_/5)^2^)^1/2^ and affected residues were mapped on the surface of eIF5-CTD, for which the backbone NMR resonance assignments were available (Paul et al., 2022). For statistical analysis, average chemical shift changes and standard deviations were calculated in Excel. For K_D_ determination, ^15^N/^2^H-labeled or ^15^N-labeled protein samples were titrated with increasing concentrations of an unlabeled binding partner, until saturation or until solubility limit was reached. Data analysis was done in SigmaPlot, as described previously (Paul et al., 2022).

k_off_ estimation from NMR data was based on its effect on exchange broadening (reviewed in (Marintchev et al., 2007)). In **Fig 3A**, peaks that move by up to 50 Hz (0.1 ppm for ^1^H on a 500 MHz NMR spectrometer) are in fast exchange (where peaks move from their position in the free labeled protein to their position in the complex with the ligand as a function of the fraction of bound protein), with no significant exchange broadening. Peaks that move by > 100 Hz (0.2 ppm for ^1^H) undergo exchange broadening, sometimes beyond detection, at ligand concentrations where a fraction of the protein is bound. Under fast exchange conditions, the line broadening δν, due to exchange is dependent on the chemical shift change between the free and bound state, Δ [s^-1^]. At equal populations of free and bound states, δν is approximately equal to (πΔ^2^)/4k_off_. Therefore, under the experimental conditions (150 mM KCl and 25 LJC), if exchange broadening becomes significant for Δ > 100 s^-1^, k_off_ is in the order of 100 s^-1^.

### Size exclusion chromatography (SEC)

SEC experiments were performed on a Bio-Rad NGC chromatography system, using Enrich SEC70 column, in buffer containing 10 mM Na Phosphate, pH 7.0, 150 mM KCl, 1 mM EDTA, 0.02% NaN_3_, 1 mM DTT and 0.1 mM AEBSF. SEC molecular weight standards (Bio-Rad) were used for calibration.

### Sequence and structure analysis

Protein BLAST (http://blast.ncbi.nlm.nih.gov/Blast.cgi) was used to obtain initial sequence alignment between human and *S. cerevisiae* eIF2β. Aligning all six K-boxes was done using Clustal O from the Max-Planck Institute for Developmental Biology Bioinformatics Toolkit (http://toolkit.tuebingen.mpg.de/), followed by manual editing. Instead of by conservation, the aminoacids were colored by hydrophobicity (green), negative charge (red), and positive charge (blue). MOLMOL (Koradi et al., 1996) was used to align the structures of human and *S. cerevisiae* eIF5-CTD, and for structure visualization.

## Data Availability

The atomic model generated in this study has been deposited in the Protein Data Bank (PDB: 9F79). The data that support this study are available from corresponding authors upon reasonable request.

## Acknowledgments

The authors thank Anvesh Sharma for assistance with NMR data analysis. This work was supported by the National Institute of Health [GM134113 to A.M.].

## Author Contributions

Conceptualization, B.K, R.F., and A.M.; Methodology, B.K, R.F., and A.M.; Investigation, P.A.W., M.S., B.K., R.F., and A.M.; Writing – Original Draft, P.A.W., B.K., R.F., and A.M.; Writing – Review & Editing, P.A.W., B.K., R.F., and A.M.; Funding Acquisition, B.K, R.F., and A.M.; Supervision, B.K, R.F., and A.M.

## Expanded View Figure legends

**Figure EV1.The eIF5 DWEAR motif contacts the eIF5-CTD.**

(A) The asymmetric unit of the crystal structure of eIF5-CTD•eIF2β-K23 complex. The two eIF5- CTD molecules are colored in yellow and blue, respectively, and the eIF2β-K23 is shown in gray. In both eIF5-CTDs, the DWEAR-motif overlaps with the loop connecting helices α1 and α2 in the N-terminal extension and engages in crystallographic contacts with a groove formed by helices α3, α5, and α8 on the neighboring eIF5-CTD molecule.

(B) Details of the interactions between the DWEAR-motif of one eIF5-CTD molecule (Mol B) and the binding pocket on the neighboring eIF5-CTD molecule (Mol A). The DWEAR-motif of Mol A points away from its globular domain and inserts into the corresponding binding pocket on Mol B. Mol B is shown in cartoon representation; Mol A is shown in surface representation with electrostatic surface charge distribution.

**Figure EV2. NMR CSP effects in human eIF5-CTD upon binding to eIF2β K-boxes.**

(A) Constructs used in this work. Folded domains are shown as boxes. Intrinsically disordered segments are shown as lines. K-boxes K1, K2, and K3 are marked and colored dark blue, blue, and light blue, respectively. The K3 segment of S. cerevisiae eIF2β visible in the crystal structure, including the αN and αC helices (**Fig 1**) is marked with a solid line. The corresponding segments surrounding the other K-boxes, corresponding to the αN and αC helices are marked with dashed lines. The four point mutations in the putative αN helix are shown with red “X”. eIF2β-NTT is blue. The eIF2γ-binding helix is orange. The homologous C-terminal HEAT (W2) domains of eIF5, eIF2Bε, and 5MP1 are orange. The DWEAR motif in eIF5 is labeled with “DW”. BHD, β-helix domain.

(B) NMR chemical shift perturbation (CSP) effects in human eIF5-CTD upon binding to eIF2β K-boxes 1, 2, and 3. Amino-acids that could not be analyzed are shown as light grey bars. Average CSP effect, average plus 5 standard deviations (SD), and average plus five SD are marked with dashed lines and labeled.

**Figure EV3. Effects of mutations in human eIF2β K-boxes corresponding to αN and αC in binding to eIF5-CTD.**

(A) NMR spectra overlay of ^15^N/^2^H-labeled human eIF5-CTD, free (black) and in the presence of 25 μM WT K1 (red), or 36 μM K1 with mutant αN helix (blue). Note that the mutant K1 causes smaller, and sometimes different CSP effects. Examples of affected peaks are boxed.

(B) NMR spectra overlay of ^15^N/^2^H-labeled human eIF5-CTD, free (black) and in the presence of 25 μM WT K3 (red), or 25 μM K3 with deletion in the αC helix (blue). Note that the mutant K3 causes smaller CSP effects. Examples of affected peaks are boxed.

**Figure EV4. eIF2Bε-CTD and 5MP1-CTD form similar complexes with each of the three K-boxes of eIF2β.**

(A, B) NMR spectra overlay of ^15^N/^2^H-labeled human eIF2Bε-CTD (A) and 5MP1-CTD (B), free (black) and in the presence of K123 (red), K1 (blue), K2 (green), and K3 (orange). Examples of peaks that are in similar or different positions in the different complexes are boxed.

(C) NMR spectra overlay of titration of ^15^N/^2^H-labeled 5-CTD with K1, with insets for a couple of moving peaks in fast and intermediate exchange.

**Figure EV5. eIF2β can bind eIF5, eIF2Bε, and 5MP1 simultaneously.**

(A) NMR spectra overlay of ^15^N/^2^H-labeled human GB1-tagged eIF2β-K123, 40 μM, free (black) and in the presence of 150 μM 5MP1-CTD (red), 150 μM eIF5-CTD (blue), 150 μM eIF2Bε-CTD (green), or in the presence of all three proteins at 70 μM each (orange). Two of the peaks affected by binding are zoomed in the inset boxes. Note that in the presence of all three binding partners, the positions of the affected eIF2β peaks are the weighed average of their positions in the presence of the individual proteins. Therefore, at the NMR time scale, binding to all three proteins is in fast exchange on the NMR time scale (each binding site on eIF2β binds to, and dissociates from each of the three proteins multiple times per second). If the dissociation rates of the individual interactions were slower, the affected eIF2β peaks would have been split into three peaks, corresponding to the three individual complexes.

(B) Overlay of SEC traces of free GB1-tagged eIF2β-K12, eIF5-CTD, eIF2Bε-CTD, and a mix of eIF2β-K12 (containing two K-boxes), eIF5-CTD, and eIF2Bε-CTD in a 1:1:1 ratio. Most of the proteins migrate as a trimeric complex; therefore, most eIF2β-K12 molecules must be bound to two eIF5-CTD and/or eIF2Bε-CTD molecules. The mix of eIF2β-K1m-K2 (where K1 is mutated) and eIF2Bε-CTD in a 1:1 ratio (red trace) served as a reference for the mobility of the heterodimeric complex. The positions of molecular weight markers are shown with vertical dashed lines.

