## Supplementary figures and images for "Molecular basis for the interactions of eIF2β with eIF5, eIF2B, and 5MP1 and their regulation by CK2"

### Supplemental Figure 1

**Figure EV1**

**A**

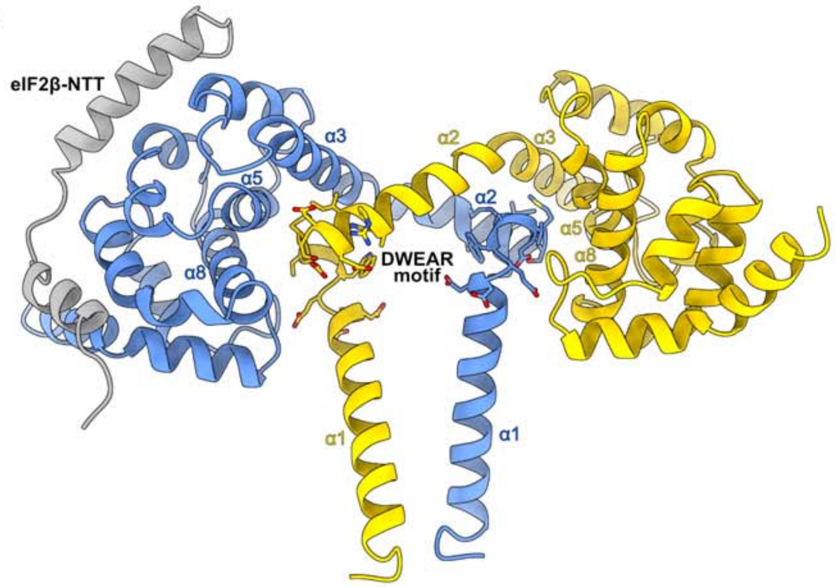


**B**

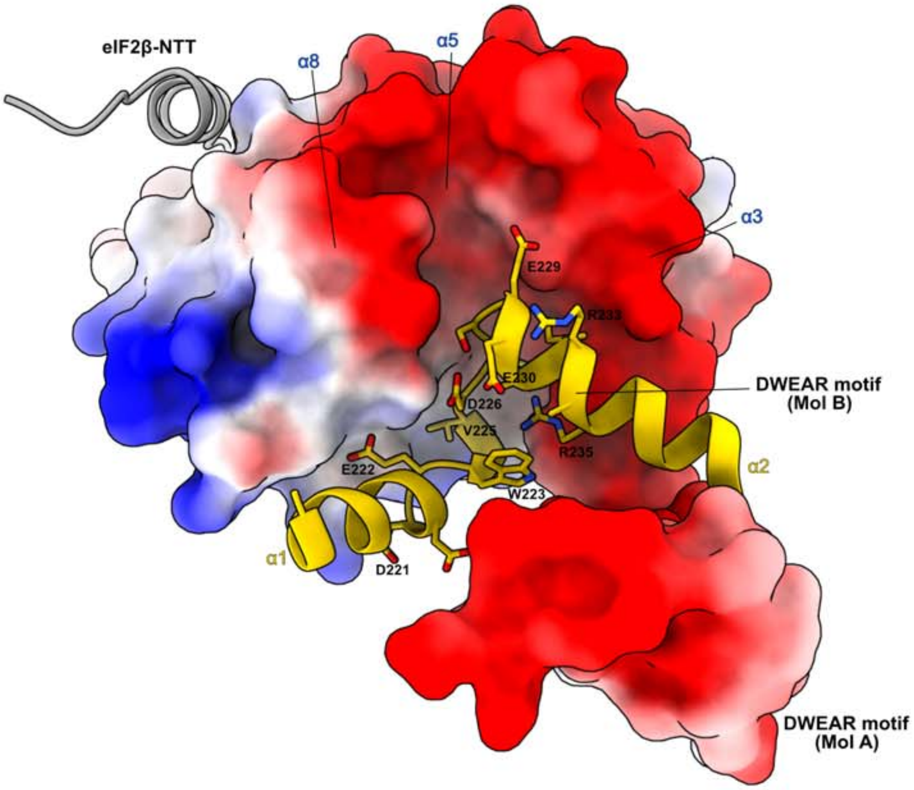
