## Supplemental Figure 2 for "Molecular basis for the interactions of eIF2β with eIF5, eIF2B, and 5MP1 and their regulation by CK2"

### Figure EV2

#### A Constructs used in this study

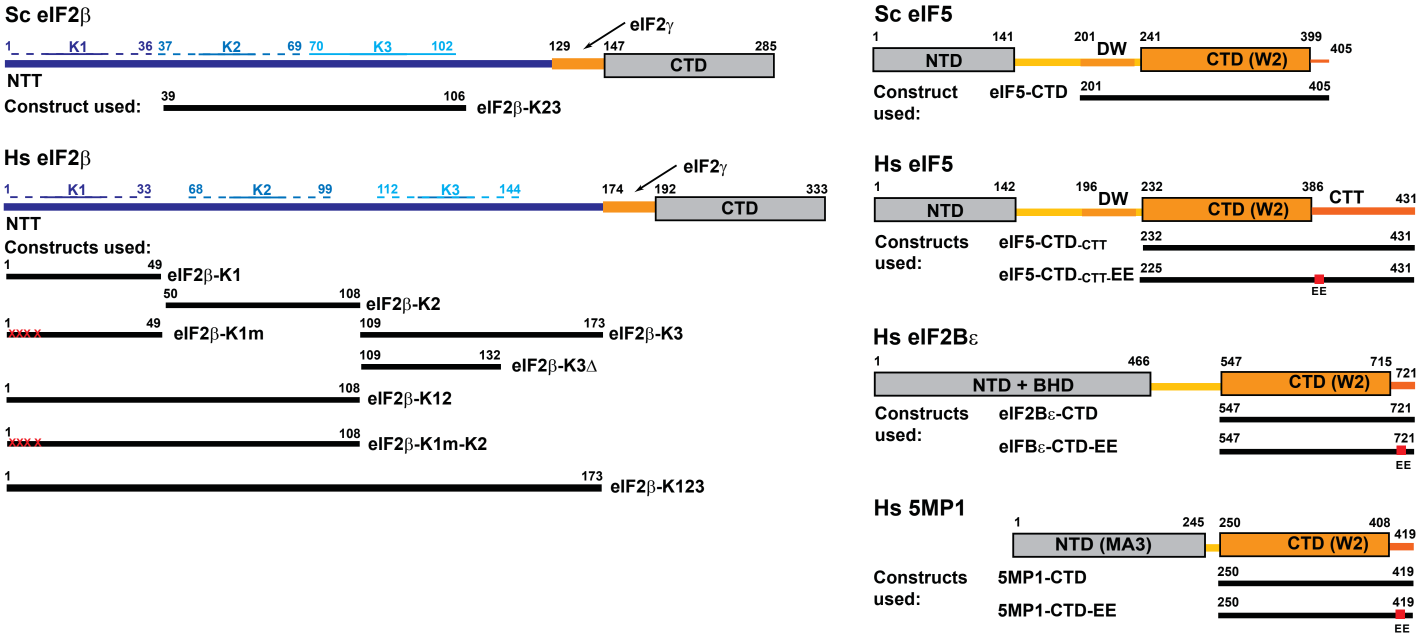


#### B NMR CSP effects in human eIF5-CTD upon binding to eIF2β K-boxes

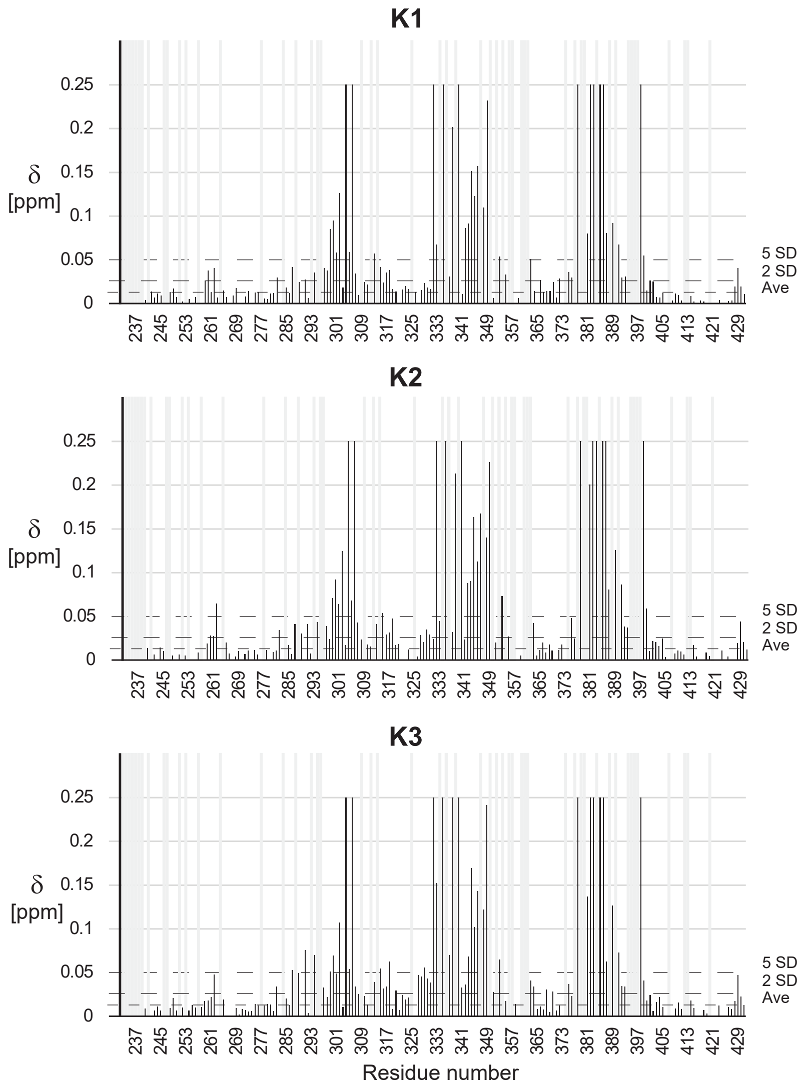
