## Supplemental Figure 3 for "Molecular basis for the interactions of eIF2β with eIF5, eIF2B, and 5MP1 and their regulation by CK2"

### Figure EV3

**A** **Overlay of NMR spectra of eIF5-CTD,50 μM, in the absence and presence of WT eIF2β K1, 25 μM, (red) and with mutations in helix αN, 36 μM, (blue)**

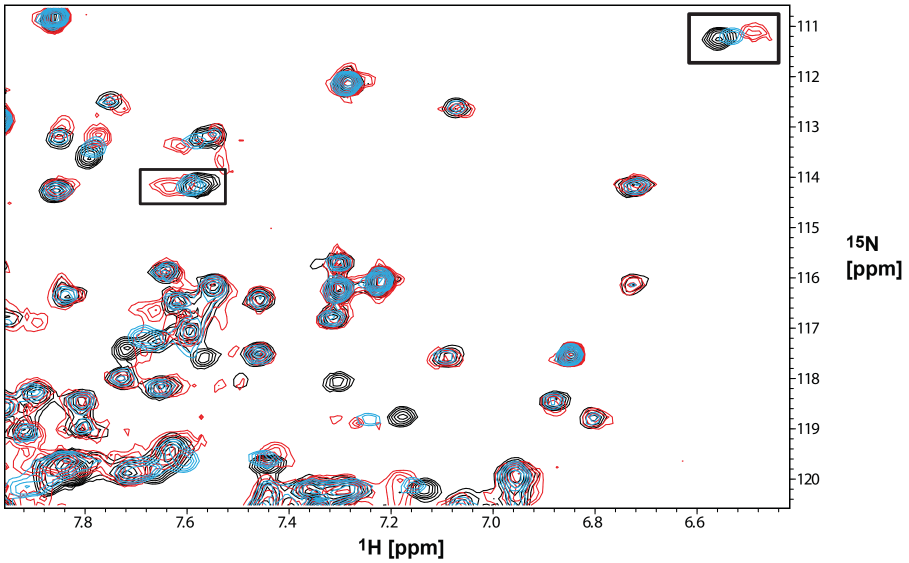


**B** **Overlay of NMR spectra of eIF5-CTD, 50 μM, in the absence and presence of WT eIF2β K3, 25 μM, (red) and with deletion in helix αC, 25 μM, (blue)**

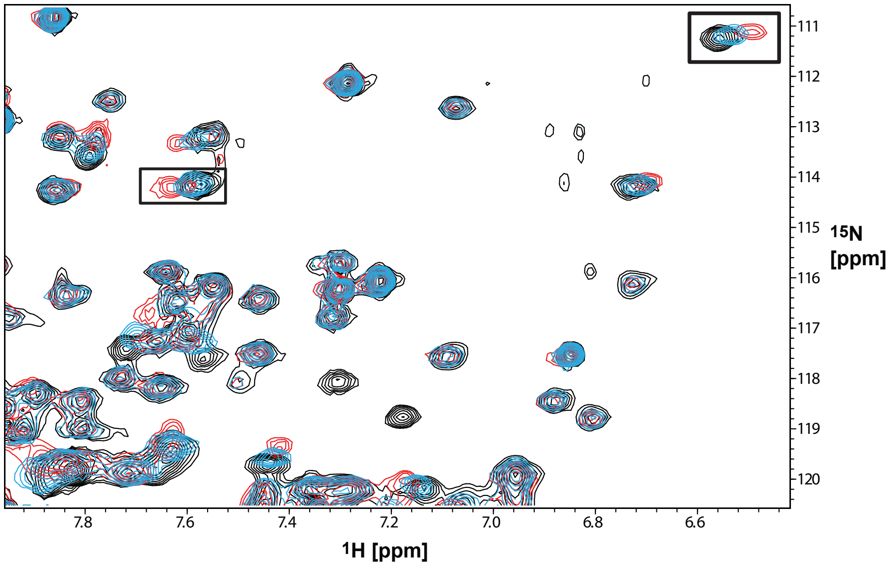
