## Supplemental Figure 4 for "Molecular basis for the interactions of eIF2β with eIF5, eIF2B, and 5MP1 and their regulation by CK2"

### Figure EV4

**A** Overlay of NMR spectra of Hs eIF2Bε-CTD in the absence and presence of different eIF2β K-boxes

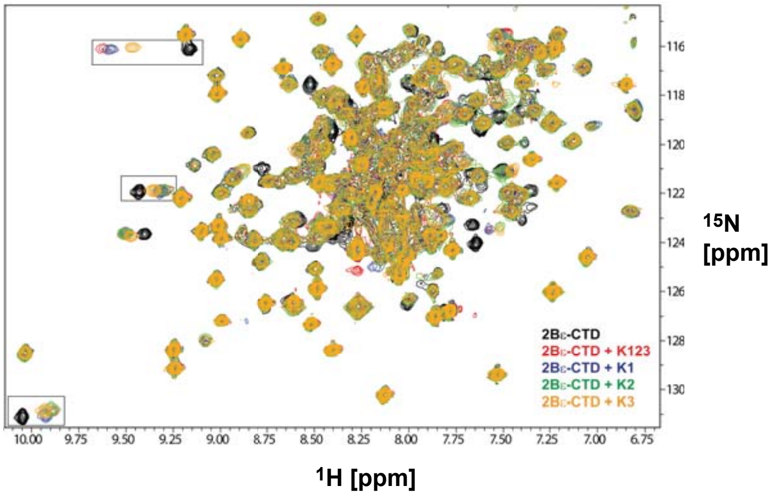


**B** Overlay of NMR spectra of Hs 5MP1-CTD in the absence and presence of different eIF2β K-boxes

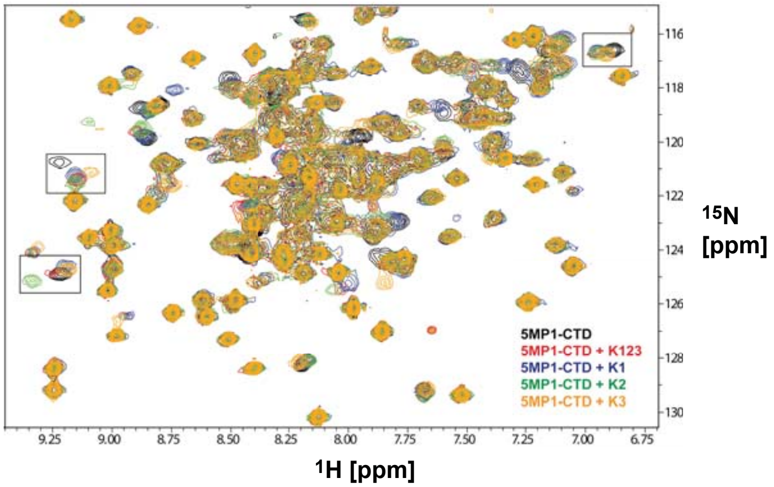


**C** NMR titration of eIF5-CTD with eIF2β K3

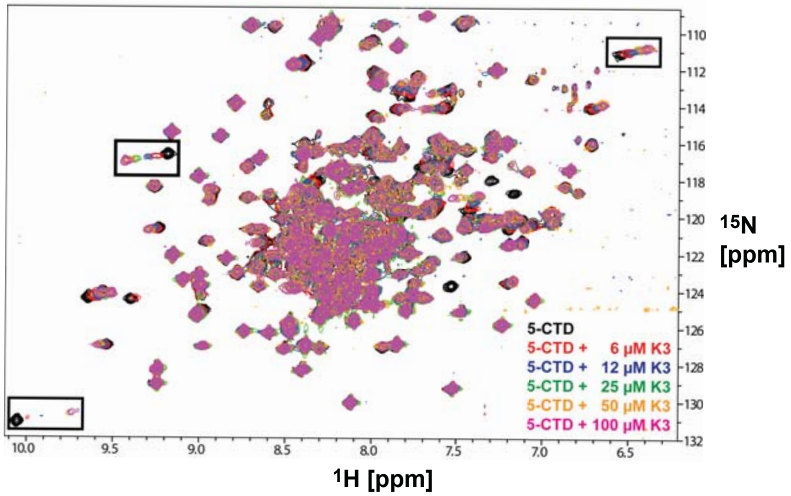
