## Supplemental Figure 5 for "Molecular basis for the interactions of eIF2β with eIF5, eIF2B, and 5MP1 and their regulation by CK2"

### Figure EV5

**A** Overlay of NMR spectra of $^{15}N/^{2}H$-labeled Hs eIF2β-K123 in the absence and presence of unlabeled eIF5-CTD, eIF2Bε-CTD, and/or 5MP1-CTD

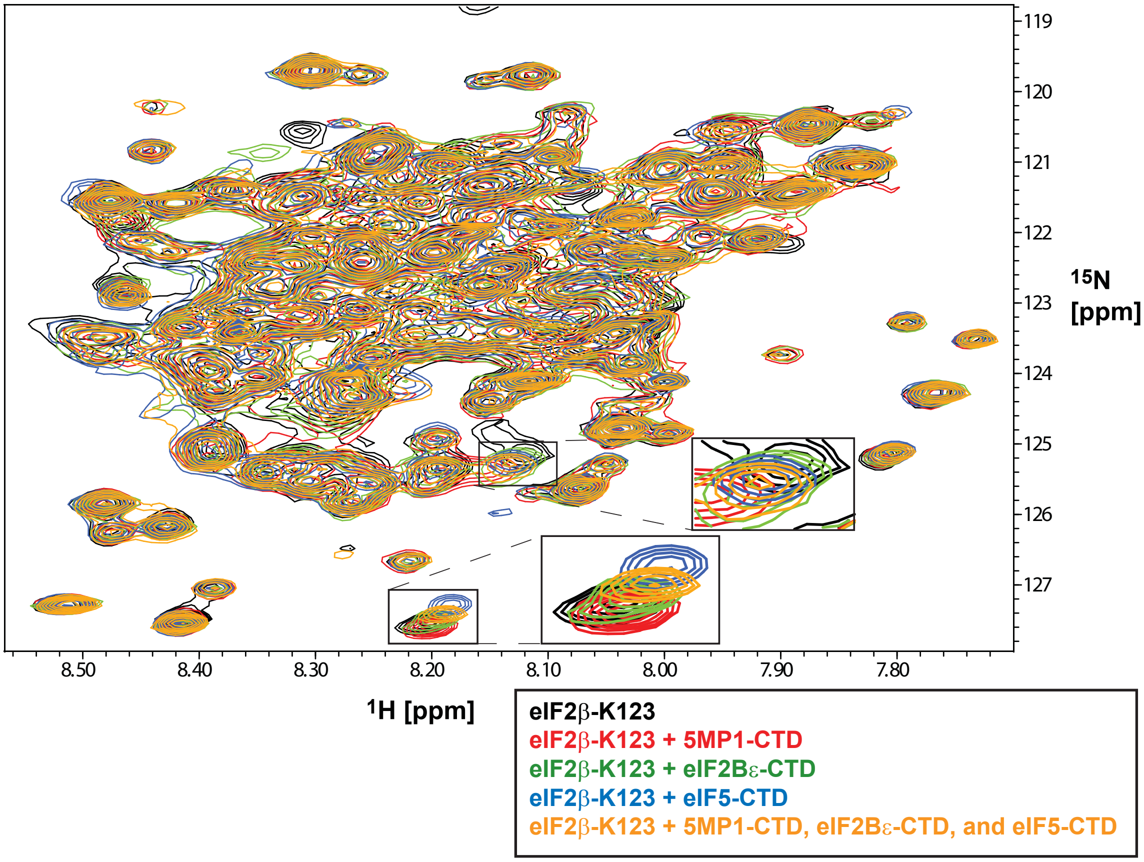


**B** Overlay of SEC traces of Hs eIF2β-K12, eIF5-CTD, eIF2Bε-CTD, and all three proteins together

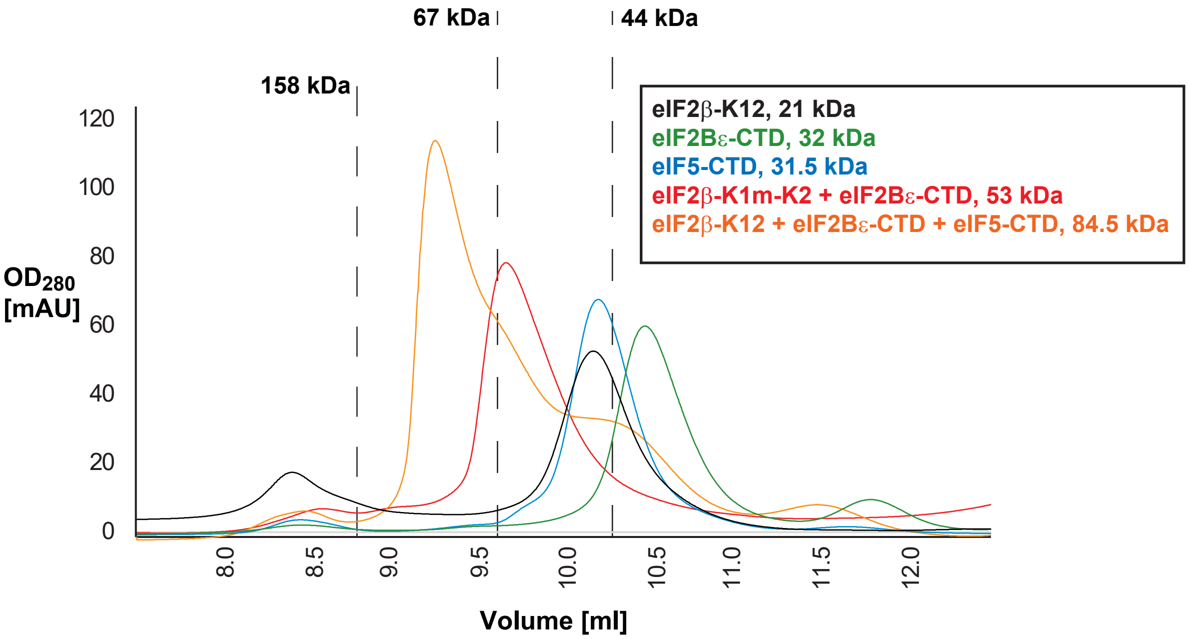
